# Distinct ground state and activated state modes of firing in forebrain neurons

**DOI:** 10.1101/2021.09.20.461152

**Authors:** Daniel Levenstein, Jonathan Gornet, Roman Huszár, Gabrielle Girardeau, Andres Grosmark, Adrien Peyrache, Yuta Senzai, Brendon O. Watson, Kenji Mizuseki, John Rinzel, György Buzsáki

**Affiliations:** Neuroscience Institute, New York University, New York, NY 10016, USA; Center for Neural Science, New York University, New York, NY 10003, USA; Department of Neurology, New York University, New York, NY 10016, USA, Langone Medical Center, New York University, New York, NY 10016, USA

## Abstract

Neuronal firing patterns have significant spatiotemporal variability with no agreed upon theoretical framework. Using a combined experimental and modeling approach, we found that spike interval statistics can be described by discrete modes of activity. Of these, a “ground state” (GS) mode of low-rate spiking is universal among forebrain excitatory neurons and characterized by irregular spiking at neuron-specific rates. In contrast, “activated state” (AS) modes consist of spiking at characteristic timescales and regularity that are specific to neuron populations in a given region and brain state. The majority of spiking is contributed by GS mode, while neurons can transiently switch to AS spiking in response to stimuli or in coordination with population activity patterns. We hypothesize that GS spiking serves to maintain a persistent backbone of neuronal activity while AS modes support communication functions.

---

A widely held view in neuroscience holds that neural computation is performed by irregular spiking that follows a smoothly-varying rate (*1, 2*). In this framework, Poisson-like spiking arises from a fluctuation-driven regime in which inhibitory and excitatory inputs are balanced (*3–7*), and maximizes the efficacy of information transmission (*1, 8*). This classical view stands in contrast to the widely-observed constraint of spike times by network oscillations (*9*) biophysical properties (*10*), and the temporally-coordinated cell assemblies that neurons form to effectively discharge their downstream target partners (*11*). Further, the statistical properties of spike patterns vary extensively between brain regions (*12, 13*) and change as a function of an animal’s behavior, sensory input, and brain state (*14–16*). Thus, the continuous irregular view maybe not be as universal as is widely accepted.

## RESULTS

To characterize the diversity of neuronal firing patterns across the brain, we examined the interspike intervals (ISIs, Fig. 1A) from six regions of the rodent forebrain during waking and sleep (see Methods; Suppl. Fig. 1). Neurons in each region had qualitatively distinct ISI distributions, with multiple modes of increased density at specific timescales (Figure 1B) that were characteristic of a given brain region and state (NREM sleep vs WAKE/REM, Suppl. Fig. 2).

**Figure 1:**
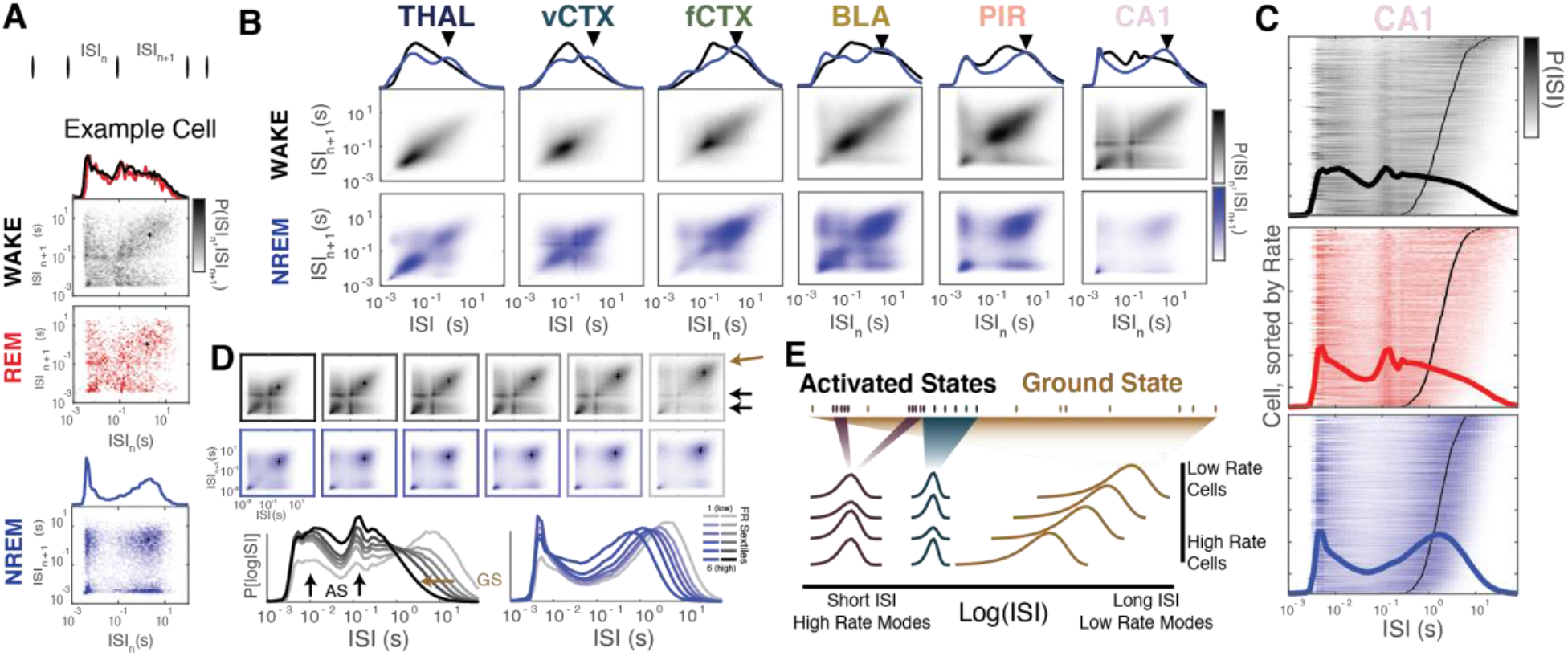
ISI distributions of forebrain excitatory neurons suggest a modal view of neural spiking with ground and activated states. **A:** logISI distribution and return map (ISI_n_ vs ISI_n+1_) from an example CA1 cell during WAKE, REM, and NREM sleep. On-diagonal clusters reflect sequential ISIs at a similar timescale, off-diagonal clusters reflect transitions between modes. **B**: Mean ISI distributions and return maps from all putative excitatory cells in each region during WAKE and NREM sleep. Black triangles reflect mean firing rate over all cells in the region/state. Due to WAKE-REM similarity, only WAKE and NREM are shown - REM can be found in supplemental figures. **C**: logISI distributions for all CA1 pyramidal cells during WAKE and NREM sleep, sorted by mean firing rate. Black line indicates the mean ISI (1/mean rate) for each cell. (See Supp Fig 3 for comparable plots from all regions). **D**: Mean ISI distribution and return maps for CA1 cells divided into firing rate sextile groups. Location (i.e. rate) of the low rate mode (ground state, GS, brown) moves to shorter ISIs in higher firing rate groups, while pattern and timescale of higher rate modes (activated states, AS, black) are consistent across firing rate group. **E**: The ground state/activated state model of neural activity.

### Ground state and activated state modes of interspike intervals

Despite heterogeneity across brain regions, we noticed a universal pattern of ISIs in excitatory neurons (Fig. 1C,D, Suppl. Fig. 2). At long ISIs (low rate), distributions tended to show a single mode that was unique to each neuron. This low rate activity corresponded to an “on-diagonal” mode in the ISI return maps, and thus reflected sequential long ISIs rather than silent intervals between episodes of higher-rate spiking. In contrast, higher-rate modes were common among neurons within each region. Each high-rate mode tended to have a distinct degree of regularity, as measured by the ISI-conditioned coefficient of variation, CV2 (Suppl Fig. 3, with higher rate modes tending to have more regular spiking (CV2 < 1), compared to those with long ISIs (CV2 ≥ 1).

These observations prompted us to consider that the distinct modes of ISI distributions are the spiking correlate of distinct neuronal and network mechanisms (Fig. 1E). We hypothesized two main categories: a ‘ground state’ (GS) mode of irregular spiking at a cell-specific low rate, and a repertoire of state-specific ‘activated state’ (AS) modes common to neurons in a given brain region.

To formalize our GS-AS hypothesis, we modeled the log-ISI distribution from each neuron with a mixture model,

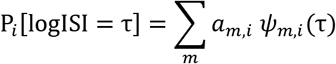

where *a_m,i_* is the weight, or fraction of spikes, from neuron *i* in mode *m*, and each mode, *ψ_m,i_*(τ), was taken to be a log-gamma distribution (*17*)(see Methods; Figure 2A-B, Suppl. Fig. 4). Fitting ISI distributions to this model allowed us to parameterize each mode by a characteristic rate (the inverse of the mean ISI) and variability (CV) which, when combined with the weights in each mode, revealed a “modal fingerprint” of the activity patterns for each neuron (Fig. 2B). In every brain region, two distinct clusters of spiking modes were seen: a low rate cluster of supra-Poisson spiking (CV>1) that corresponded to the GS mode in each cell, and a cluster of sub-Poisson modes at higher spike rates with characteristic sub-clusters that were unique to each region and state (Fig. 2C). To further characterize the repertoire of AS modes in each region, we constrained the model such that the properties of each AS mode were shared across neurons in the same region/state (Fig 2C, Suppl. Fig. 5, Methods).

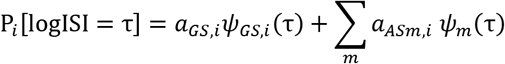

**Figure 2:**
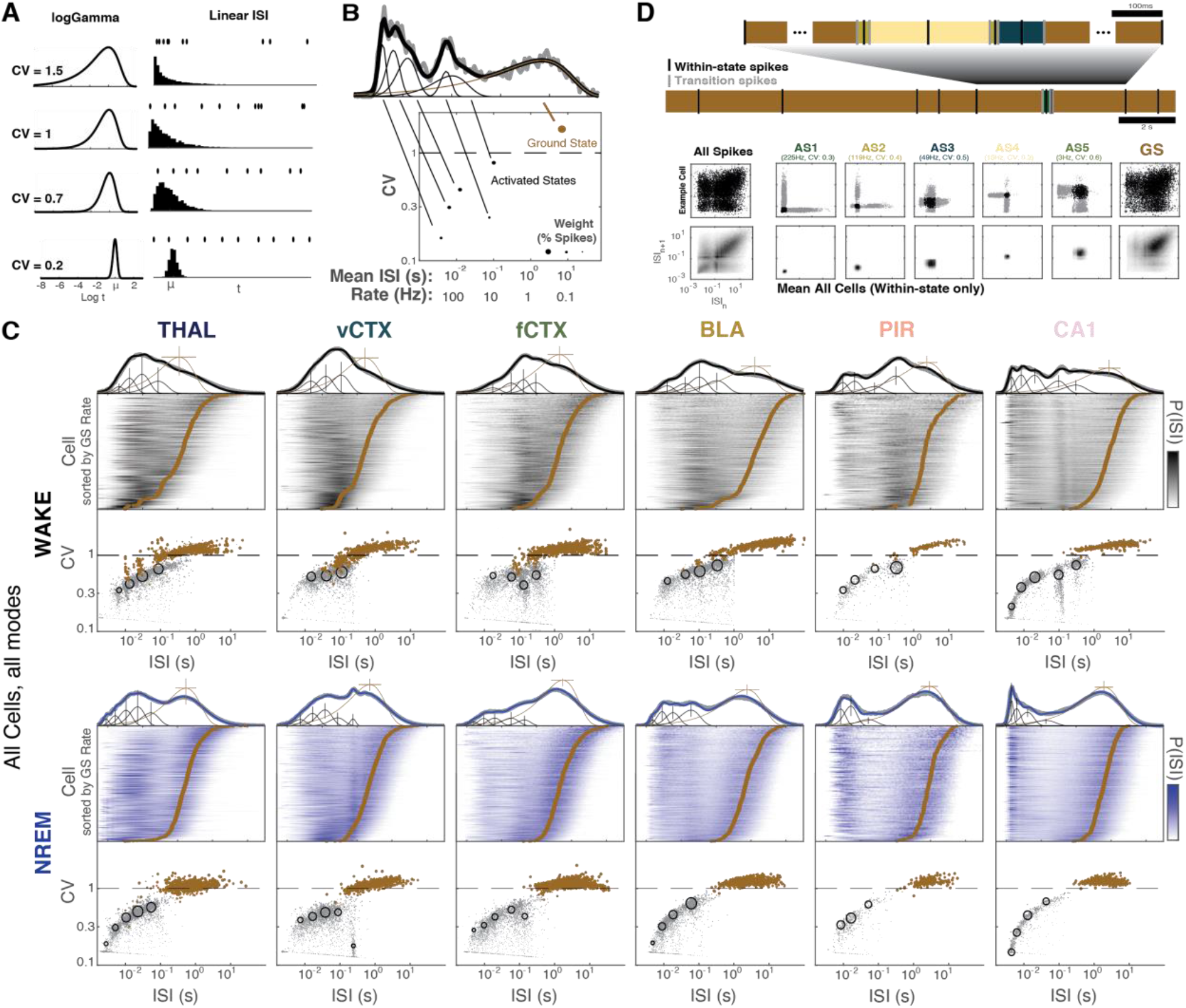
Modal decomposition of ISI distributions and spike trains. **A:** Log-Gamma distribution as a model for (log)ISI modes. A log-gamma distribution is characterized by a mean interval (i.e. a mean rate) and coefficient of variation (CV), which reflects the irregularity of spiking in that mode and can range from regular (CV<<1), to Poisson-like (CV~1), to supra-Poisson (CV>1) spiking. Four examples of spike trains and ISI distributions sampled from four log-gamma distributions with a range of CVs are shown, each with mean ISI = μ. **B**: Mixture of log-gammas model fit for an example CA1 cell during WAKE. The logISI distribution for the cell is decomposed into a sum of log-gamma distributions. Each mode is represented by its mean ISI (inverse of the mean rate) and CV, and a weight, which reflects the proportion of spikes fired by the cell in that mode. The example cell has a ground state at 0.1Hz with a CV=2, and 5 activated state modes at different rates and CVs. **C**: Log-gamma mixture model fit for all cells in each region/state. Grey points correspond to individual cell fits. Open circles correspond to model fit with each AS mode constrained to have the same mean/CV over all cells (”shared” AS modes, see Methods), sized by the mean weight over all cells. **D**: Hidden Markov Model for decomposing spike trains into modes.

We then used a hidden Markov model (HMM) approach to assign each ISI to its most likely mode (Methods, Fig. 2D).

Many AS modes corresponded to known activity patterns in each region. For example, CA1 neurons during WAKE had an AS mode with regular spiking ISIs at ~100 ms that was associated with theta oscillations (~9Hz) (Supp Fig 6, (*18*)), and vCTX neurons had an AS mode of regular spiking at ~250ms which was associated with delta oscillations (Supp Fig 6 (*19*)). Further, neurons in each region had prominent AS modes at gamma oscillation timescales of 10-40 ms that were often associated with local interneuron activation (*20*)(Supp Fig 7). Finally, a “burst” mode of regular spiking at <10 ms was most prominent in the hippocampus during NREM sleep (*21–23*), (Suppl. Fig. 8).

### Activated states of spiking

To examine conditions that activate neurons out of the ground state, we examined ISI statistics during specific behaviors. The spiking of a subset of hippocampal neurons increased at specific locations, consistent with known place field activation (*24*). When averaged across trials, spike rate varied smoothly with position and peaked at specific positions in the environment with different peak rates. However, when we plotted ISIs as a function of position (Fig. 3A), this increased rate of spiking corresponded to the appearance of a discrete multi-modal cloud of 10-150ms ISIs, which was similar among all place cells (Fig. 3B-D). When we compared the in-field and out-field ISI distributions and return maps, we found that they differed in the relative occupancy in high- and low-rate modes, rather than reflecting a qualitative shift in the shape of the distribution (Fig. 3E). This change in mode occupancy reflected decreased incidence of GS spiking and increased incidence of 100ms, 20ms, and 8ms AS modes within the place field, as identified by the HMM (Fig. 3B) and quantified by changes in mode weights when in-field and out-field ISI distributions were compared with the mixture-of-log-Gammas model (Fig. 3F; Methods).

**Figure 3:**
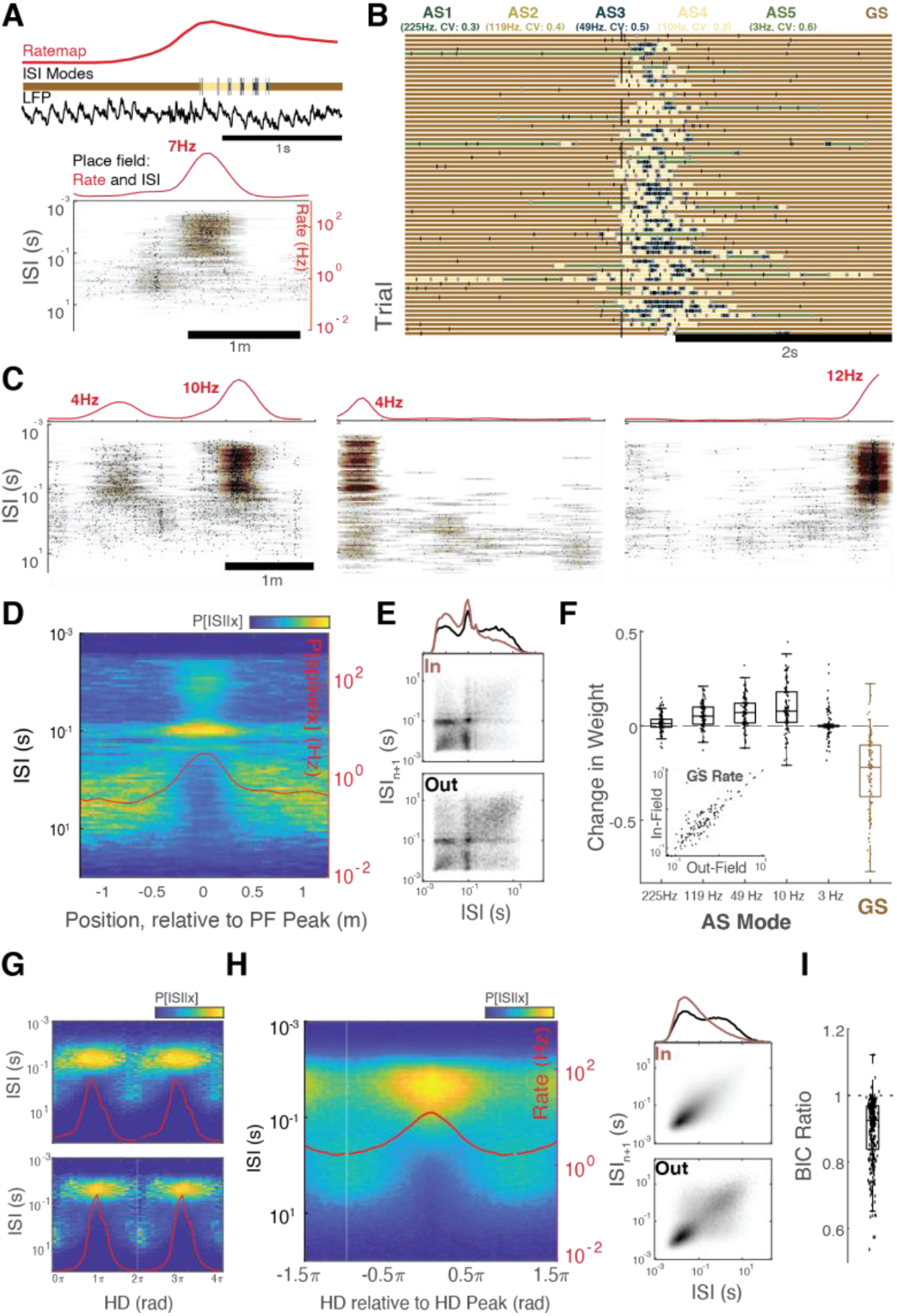
External inputs evoke activated states in hippocampal place cells and thalamic head direction neurons. **A:** An example place cell. (Top) LFP, spikes, and spiking modes for a single traversal of the place field. (Bottom) adjacent ISIs from all spikes from all traversals, as a function of position. **B:** Trial-by-trial spiking modes, as classified by the HMM, for the example cell in A, aligned to entering the place field. **C:** Additional three example place cells. Despite variation in peak rate, in-field spikes have ISIs of similar timescales. **D:** Average position-conditional rate (P[spike | position; red line]) and ISI distributions (P[ISI | position]) for CA1 place cells, centered on their place field peak. **E:** Mean in- and out-field ISI distributions and ISI return maps. In- vs out-field ISI distributions do not differ in the general shape of the ISI distribution, but in the relative occupancy in activated state and ground state modes. **F:** Place fields increase occupancy of theta and gamma-timescale spiking modes, and decrease occupancy of GS modes. Change in weight for each mode between in- and out-field ISI distribution, as measured by the mixture of gamma model, for all CA1 place cells. (Inset) GS rate remains similar in and out of the place field. **G:** Rate (P[spike | HD], red line) and ISI distribution as a function of head direction (HD) for two cells in the AD thalamus. **H:** Average rate and conditional ISI for all HD cells, centered on their preferred head direction. While rate varies continuously with position and head direction (red line), ISI distribution shows distinct modes of activity as a function of HD. (Right) Mean in- and out-field ISI distributions and ISI return maps. **I:** Modal encoding model fits HD cell activity better than continuous rate encoding model (Supp Fig 9). Bayesian information criterion (BIC) ratio for model fits from each HD cell; ratio <1 indicates better fit to the modal model.

To examine the generality of these findings, we analyzed thalamic head direction neurons (*25*) in a similar manner. Smooth variation in cells’ average firing rate as a function of head direction masked underlying discrete ISI modes. The timescale of ISIs in each mode was constant regardless of head direction, but occupancy was highest in the AS mode when the rat was gazing at the preferred direction of the neuron, whereas GS spikes dominated the less preferred directions (Fig. 3G, H). Because of the unimodal nature of the activated states in head direction neurons, we were able to directly compare a “continuous rate” encoding model with a discrete “modal” encoding model (Methods, Fig. Suppl Fig. 9),

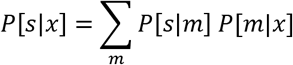

in which the probability of observing *s* spikes in a given time window is determined by the head direction-dependent probability of a cell being in mode *m* (P[*m*|*x*]) times the probability of observing *s* spikes in that mode (P[*s*|*m*]). The modal model overwhelmingly fit the observed spiking data better than the continuous rate model (Fig. 3I), further supporting the modal view of spiking.

### Ground State spiking reflects a default mode of balanced activity

In contrast to region/state-specific AS modes, GS mode (Fig. 4A) was present in every brain region during all brain states. A neuron’s mean firing rate was more strongly correlated with its GS rate than with the occupancy of its AS modes or their rate (Fig. 4B,C), mainly because GS spikes represented the majority of spikes in most cells (Fig. 4E), and the vast majority of time was spent in the long-duration intervals between GS spikes (Suppl Fig. 10). Similarly, a neuron’s GS rate was more strongly conserved across WAKE and NREM states compared to changes in the occupancy of AS modes (Fig. 4F).

**Figure 4:**
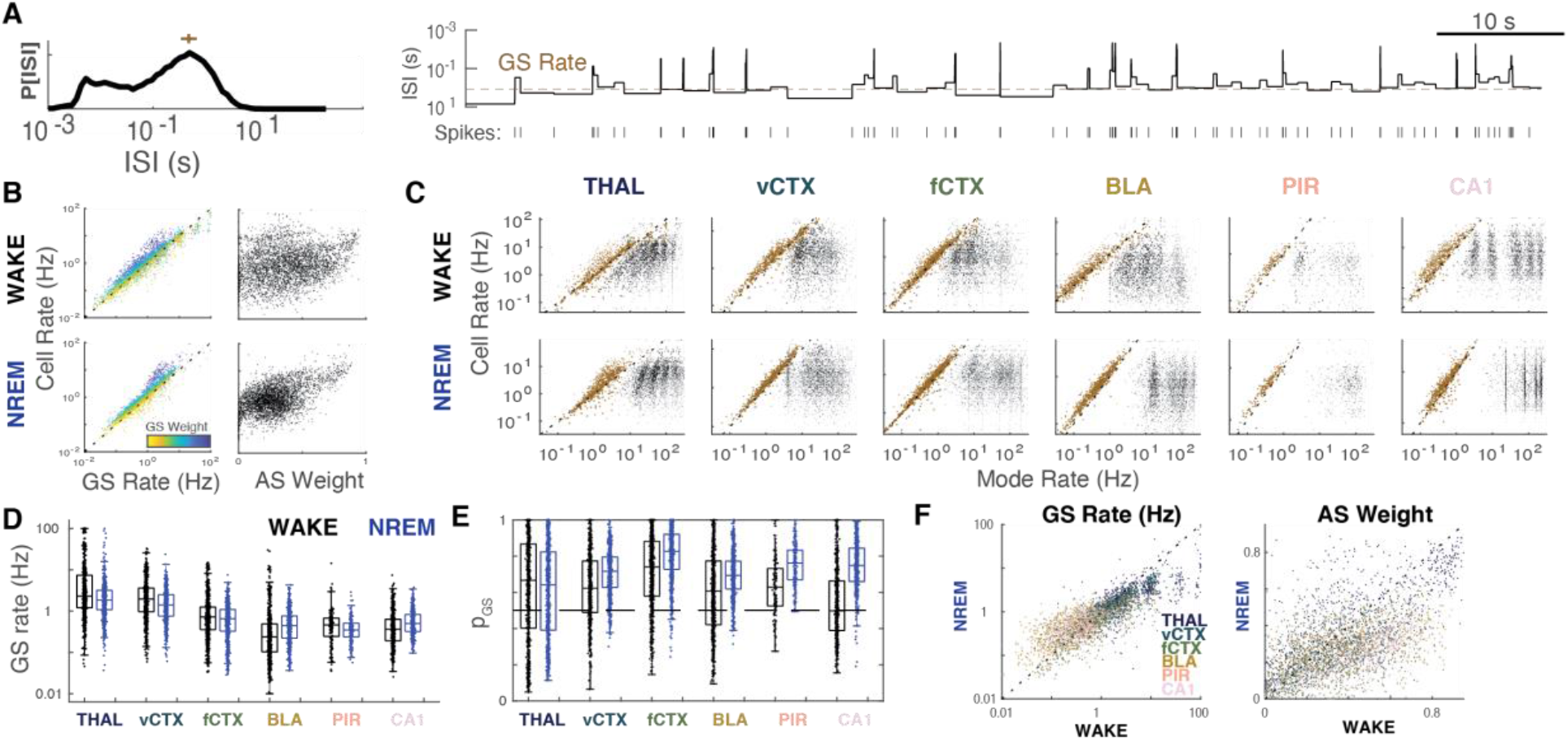
The Neuronal Ground State. **A:** Ground state activity in an example hippocampal neuron - NREM. Left, ISI distribution. Cross indicates ground state rate. Right, sample spike train and ISI from the example cell. Note that the majority of time is spent in intervals around the GS rate (dashed line). **B**: Mean rate as a function of GS rate (left) and total AS weight (right) in all cells in all regions during WAKE and NREM. Note: AS weight is equivalent to 1-GS weight **C**: Mean rate as a function of modal rate from all modes from individual fits of all cells. Each cell can have up to 6 points in each plot, reflecting GS rate and the rate of up to 5 AS modes. In each region, rate of the GS mode (brown) is tightly correlated with mean rate, unlike AS modes. **D**: GS rate for all cells. **E**: GS weight (p_GS_, fraction of spikes in the ground state) for all cells. GS spikes compose more than 50% of the ISI distribution in most cells from all regions/states. **F**: Comparison of GS rate and total AS weight of all cells in WAKE and NREM.

Due to its irregular spiking (CV≥1), we hypothesized that the GS mode arises from balanced inhibition and excitation (*4*, *7*). We tested this idea with a network model of integrate and fire neurons (*26*) in which each neuron used an inhibitory plasticity rule (*27*)(Fig 5A) to maintain a cell-specific target rate, which were lognormally-distributed across the population (Fig 6B, Methods). Under conditions of moderate excitatory (E-E) recurrent connectivity and near-threshold drive, the network showed a balanced regime of asynchronous activity (*26*), in which each excitatory neuron spiked irregularly at a cell-specific rate akin to GS-like activity (Fig. 5B,C). When we varied the strength of recurrent excitation or external drive, additional activity patterns were induced in the network, including E/I (“gamma”-like) oscillations (*20*), a heterogeneous asynchronous regime (*28*) and network bursts (Fig. 5D, E). Each of these regimes produced a characteristic pattern of AS modes in the ISI distributions that was common to all neurons in the network and reflected their engagement in collective activity. However, in each regime, GS activity was also preserved in the spiking of single neurons despite different collective dynamics.

**Figure 5:**
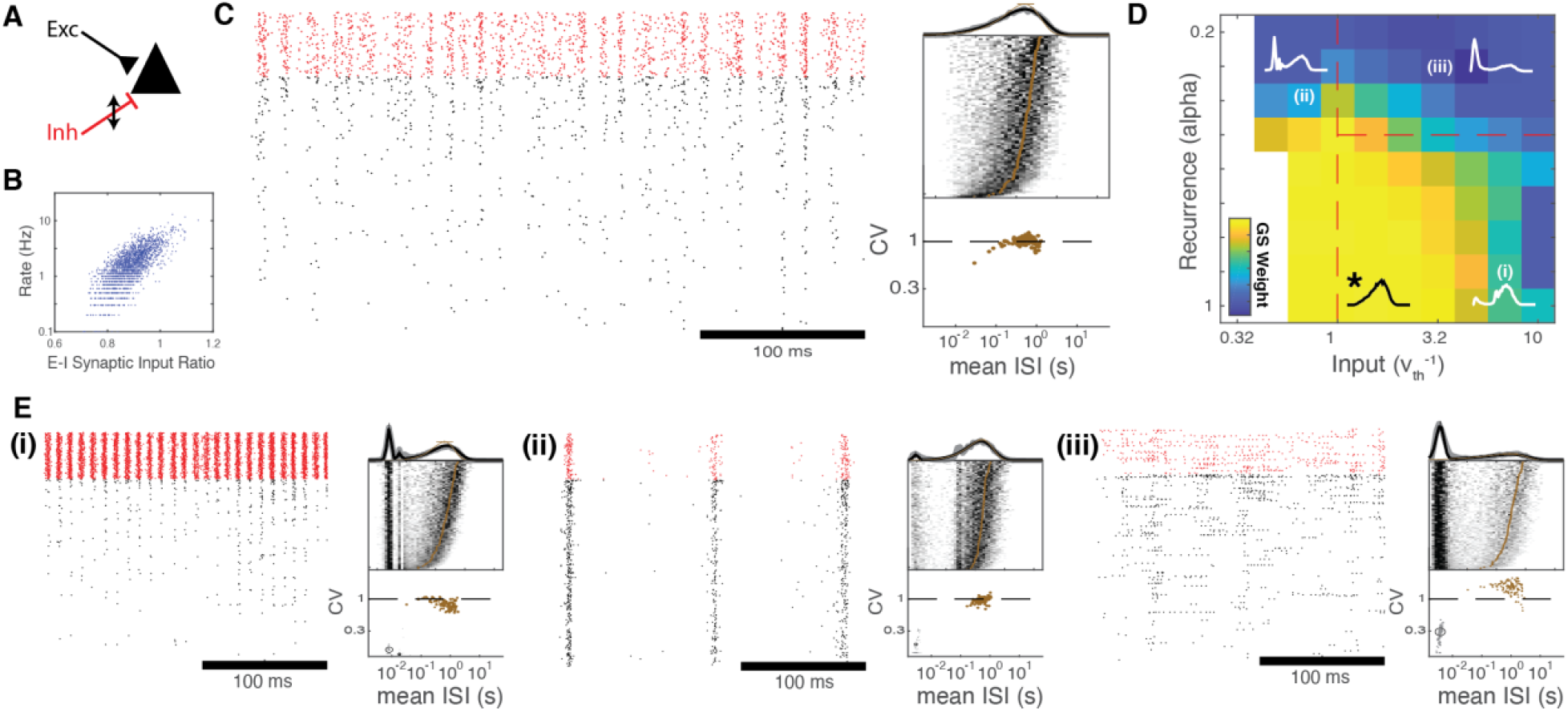
GS and AS ISIs are present in a heterogeneous self-balancing network. **A**: The heterogeneous self-balancing network. Each cell maintains spiking at a cell-autonomous rate by adjusting its incoming inhibitory synapses to balance excitatory inputs. **B**: GS rate reflects E/I synaptic input ratio. Firing rate is determined by the ratio of incoming excitatory and inhibitory synaptic weights in each cell. **C**: Left: Spike raster from excitatory (black) and inhibitory (red) cells. Right: ISI distribution of a subset of cells in the network, sorted by GS rate, as identified by the mixture of gammas model. **D**: Mean GS weight over cells in the network, as determined by the mixture model, as a function of E-E synaptic weights (recurrence) and the level of external drive. External drive and increased recurrence reduce GS weight by producing network activation patterns, each with associated AS modes (panel E), but GS mode is maintained. Recurrence is in units of K^-alpha^, where higher values of alpha signify lower excitatory weight. **E**: Simulated rasters (left) and ISI distributions from a subset of model neurons fitted to the mixture of gamma model (right) for neurons in self-balancing networks under conditions of strong drive (i), strong recurrence with low drive (ii), and strong recurrence (iii).

## DISCUSSION

Most previous research has focused on the question of how neurons respond to sensory stimuli or induce motor outputs by changing a continuously varying rate of Poisson-like spiking. In contrast, our results suggest a framework in which single neurons engage in distinct spiking modes. The “ground state” (GS) mode is universal among neurons in all observed regions and characterized by irregular spiking at a cell-specific low rate. In addition to the GS mode, neurons have a repertoire of higher rate “activated state” (AS) modes, with more regular spiking at characteristic timescales that are common among neurons in a given region and brain state. We hypothesize that GS mode maintains the brain’s internal dynamic, while AS modes serve communication functions.

We suggest that AS modes are the spiking correlates of brain state-specific activity patterns that emerge from the unique network and cellular properties of a given region. AS modes could often be related to known regional population patterns, as identified by LFP oscillations or tuning to external stimuli. The more regular, higher rate spiking of AS modes at particular timescales is ideal for effective transmission (*29, 30*) and multiplexing (*31, 32*) of functionally-distinct signals to downstream readers. AS spiking can exert mode-specific impact on postsynaptic cells (*33*) and networks (*34*), or activate intracellular processes in the spiking cell itself with mode-specific implications (*35–37*).

In contrast to AS modes, slow irregular spikes are often considered as “noise”, although the potentially beneficial role of noise has been acknowledged (*38*, *39*). Because of the long ISIs of GS spikes, a target pyramidal neuron with short integration time constant (<50 ms) will not be able to effectively integrate the rate of GS spikes from each of its upstream partners. Yet, given that GS spikes represent the majority of spikes in the brain, their functional significance is expected to be important (Suppl. Fig. 11). We propose that the GS mode of spiking serves the primary function of brain networks which is to maintain their own dynamics. The combined effect of GS activity might keep neurons in a near-threshold state from which they can respond quickly to relevant inputs (*39, 40*), and even solitary spikes can induce responses in target interneurons (*33*) or sparse strongly-connected pyramidal cells (*41*). This ongoing background of GS activity may maintain a diverse reservoir of activation patterns that are available to be selected and reinforced when associated with salient behaviors or stimuli (*42–45*).

## Author Contributions

DL, GB and JR conceived the experiments and framework, GG, AG, AP, YS, and BW performed the experiments, DL designed and executed all data analyses and computational modeling with assistance from RH on the HMM analysis and JG on the development of the spiking network, DL, GB and JR wrote the manuscript with contributions from other authors.

## Acknowledgements

The authors would like to thank Xiao-Jing Wang, Bijan Pesaran, Biyu He, Kenneth Harris, and Rachel Swanson for feedback on an early version of the manuscript. Supported by NIH MH122391, U19 NS104590, U19NS107616.

## Data availability

The datasets and code used in this study are available (data: https://crcns.org, code: https://github.com/buzsakilab/buzcode)

## Supplementary Materials

### Experimental data

All data used in this study were previous published (*14, 19, 25, 46, 47*) and are available at: https://buzsakilab.com/wp/database/ and on the CRCNS database (crcns.org). Units were separated into putative excitatory and inhibitory neurons in each region using the classifications specified in each paper. Our current analysis is focused on principal neurons. Limited results from interneurons are shown in Supp Fig. 9.

### Brain State Detection

The brain state scoring algorithm was based on Watson et al 2016, with modifications in Levenstein et al 2019, and with further modifications as outlined here (Supp Fig. 1). Sleep scoring was done using three main metrics: the EMG, narrowband theta power, and the slope of the (log-log) power spectrum (PSS; (*48*)). Channels used for theta and PSS were automatically selected (and manually checked) to be the channels that had the most bimodal distribution in theta power and PSS, per Hartigan’s dip test (*49*).

PSS was calculated in sliding 2-second windows (dt = 1s) using the IRASA method (Wen and Liu 2016; Supp Fig. 1A) and then smoothed in a sliding 10s window. To calculate PSS in the frequency range between 4 and 90 Hz, the FFT spectrogram was calculated for 336 evenly logarithmically spaced frequencies between 1.38 and 260.79Hz (resulting in 200 frequency bands between 4 and 90Hz, inclusive). For each frequency between 4 and 90Hz, the power was taken to be the median value of log(power) in all frequencies within a factor of 2.9 times larger and smaller (i.e., the 68 evenly-spaced log[frequencies] on each side), resulting in a median-smoothed spectrum that largely removed narrow-band peaks. The slope of median-smoothed spectrum in each window was then estimated using a linear fit (equivalent to the power law exponent of the linear spectrum), which was taken to be the PSS. The narrowband theta signal was taken to be the maximal difference between the non-smoothed spectrum and the median-smoothed spectrum in the range of 5-10Hz (Supp Figure 1A).

Brain states were then determined as outlined in Levenstein et al (2019; Suppl Fig 1B), by first using the bimodal PSS distribution to separate NREM (more negative PSS) from WAKE/REM (less negative PSS), and the bimodal theta and EMG distributions to divide REM (high theta/low EMG) from WAKE (everything else). Scoring was manually checked in all recordings. All regions showed comparable distributions of the state scoring metrics (Supp Figure 1C). Scoring code is available at https://github.com/buzsakilab/buzcode, using the function SleepScoreMaster.

### Mixture of Gammas Model and fitting

We modeled log ISI distributions as a mixture of log gamma distributions, in which one mode was unique to each neuron (i.e. the ground state, *ψ*_0_), and *N_AS_* gamma modes were shared across the population (i.e., activated states),

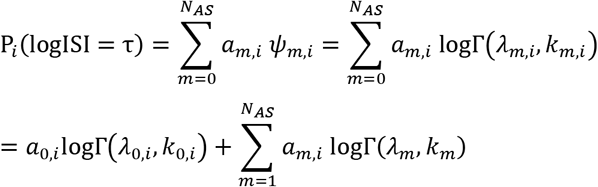

Where *P_i_*(logISI = τ) is the probability density of observing an interspike interval from cell *i* of duration *e*^τ^,

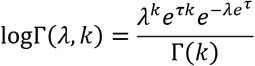

is the log-gamma distribution with total density = 1. *a_m,i_,m* ∈ {0, …, *N_AS_*}, 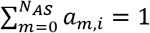 are the (non-negative) weights that correspond to the proportion of spikes from cell *i* in each mode, *λ*_0,*i*_ and *k*_0,*i*_ are the shape and scale parameters of the ground state mode for cell *i* and *λ*_*m*>0_,*k*_*m*>0_ are the shape/scale parameters for each of the *N_AS_* shared modes across the population. Throughout the text, the parameterization *rate* = 1/*kλ* and 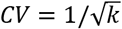 is used for readability.

To fit the model, the logISI probability density,

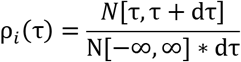

where *N*[*x,y*] is the number of ISIs between *e*^x^, and *e*^y^, was calculated for all putative excitatory cells. The ISIs from the same region/state were simultaneously fit by using the *sqp* algorithm as implemented by MATLAB’s *fmincon* function to find the parameter set *θ* ∈ {*λ*_0,*i*_,*k*_0,*i*_, *a*_0, *i*_, *λ_m_*, *k_m_*, *a_m,i_*} that minimizes the loss function:

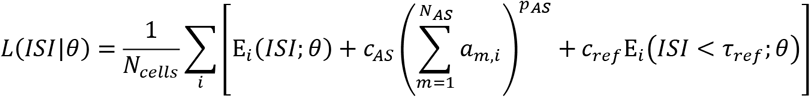

Where 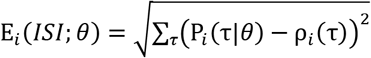 is the fitting error for cell *i*; *c_AS_* = 0.3 applies an L^1^ regularization (*p_AS_* = 1) penalty on AS mode weights to facilitate sparse fitting for each cell with a small number of modes; *c_ref_* = 2 penalizes model density for short ISIs within a refractory period *τ_ref_*. The number of AS modes, *N_AS_*, was selected by iteratively increasing the number of modes (Supp Fig 5), and was chosen based on visual inspection of the drop in *L*(*ISI*|*θ*) at each additional mode and P*_i_*(τ) such that spiking modes, when possible, matched known LFP oscillatory patterns and were able to encompass a shared mode for those seen in single cell fits. In each iteration, *λ_m_* and *k_m_* were preserved from the previous iteration, and an additional mode was initialized with *λ_m_* at the location of maximal missing ISI density and *k_m_* = 20. To acquire the single-cell fit, the cells were then re-fit without the constraint of activated states with shared shape/scale parameters, using the shared fit parameters as initial conditions.

### Modulation of spiking mode weight

To calculate the modulation of spiking mode weight by various variables (x, e.g., theta power, delta power, interneuron activity Supp. Fig 6,7), conditional ISI distributions P[ISI|x] were calculated by binning x into 8 percentile bins and calculating the distribution of ISIs adjacent to spikes that occurred in each bin. The mixture of gammas model was then fit to the ISI distributions from each bin, while constraining the AS mode rate and CV to have the same values in the full distribution fit. Modulation of each mode was taken to be the correlation between that mode’s weight and the percentile of x. For binary variables (SPW-R, within-place field, within-tuning curve; Figure 3) two bins were used (e.g., in-field vs out-field), and the modulation was taken to be the difference in weights.

### Hidden Markov model for visualization of mode sequence decoding

To assign each ISI to the most likely underlying mode, we fit a hidden Markov model (HMM) to the sequence of ISIs observed for each pyramidal neuron(*50*). The HMM assumes that each ISI is generated by a small set of latent variables *m* whose transition structure is first-order Markov: i.e., *P*(*m_j_*|*m*_*j*–1_, …, *m*_0_) = *P*(*m_j_*|*m*_*j*–1_), where *j* denotes the ordinal position of the observation (i.e., a specific ISI) in the sequence. In our treatment, *m* refers to the discrete spiking modes – a single ground state mode, and N_AS_ refers to the number of activated state modes. As described in the section above, under each mode, ISIs were assumed to follow a log gamma distribution (*17*). Furthermore, the shape/rate parameters (λ *and k*) of each mode were fixed to those discovered by the sequence-less mixture of gammas model, which leverages information across neurons when fitting parameters associated with activated state modes. Thus, the only parameters left to fit were the entries of the transition matrix *A* -- the transition probabilities (each row is a probability distribution of going from one mode to all others), and the initial state probability vector π, specifying the probability distribution of mode occupancy for the first ISI in the sequence. Prior to fitting the parameters, transition and initial state probabilities were initialized by drawing from a uniform distribution and normalizing to make values sum to 1. Parameters were then fit using the Baum-Welch algorithm (*51*). To avoid a locally optimal solution, the HMM was refit 10 times with different initializations, and the one maximizing the data log likelihood was selected. The assignment of each ISI to an underlying mode was determined via the Viterbi algorithm (*51*), which identifies the most likely sequence of modes given the observed sequence of ISIs.

### Modal and continuous-rate encoding models

We compared a continuous-rate encoding model, P[*s*|*x*] = P[*s*|*r*(*x*)], in which the probability of observing *s* spikes in a given time window is a function of a tuning curve *r*(*x*) that specifies the rate given a stimulus variable *x*, with a modal (quantal) encoding model

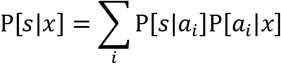

in which s, the probability of observing *s* spikes in a given time window, is determined by the spiking mode, *a_i_*, a cell is in, and the probability of being in a given spiking mode is determined by the stimulus variable, 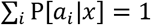. In the case of two modes *i* ∈ {*GS,AS*}, we have

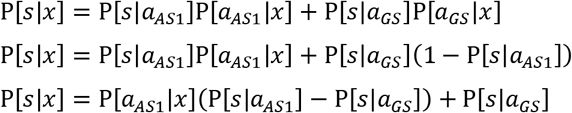

In which P[*s*|*a_GS_*] and P[*s*|*a*_*AS*1_] are the probability of observing s spikes when the cell is in GS mode and AS mode, respectively, and P[*a*_*AS*1_|*x*] is a modal “tuning curve” and specifies the probability that the cell will be in AS mode given stimulus variable x. For a circular variable (e.g., head direction), we can take P[*a*_*AS*1_|*x*] as a circular gaussian with peak of width k centered around x0

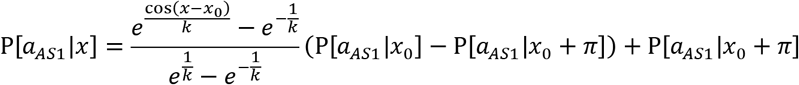

For simplicity, we take spike generation to be Poisson distributed, 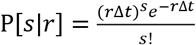, and each mode to be defined by a Poisson process with rate *r_i_*. However, this formalism could be used for an arbitrary spike count distribution P[*s*|*a_i_*], and an arbitrary number of spiking modes.

Spike count observations from head direction cells were fit to both models, using *sqp* algorithm as implemented by MATLAB’s *fmincon*, and were directly compared using the Bayesian information criterion (BIC, Figure 3I)

### Self-balancing network model

The self-balancing network was modeled using a network of leaky integrate-and-fire neurons with jump synapses as in (*26*), simulated using the Euler approximation with dt = 0.1ms.

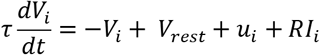

Where 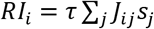 is the total synaptic input to each cell *i*, *J_ij_* is the synaptic weight from cell *j* to cell *i*, and *s_j_* is set to 1/dt following a fixed synaptic delay after each spike of cell *j*, resulting in a jump of magnitude *J_ij_* of the voltage of cell. *J_ij_* were positive for excitatory cells and negative for inhibitory cells. External input, *u_i_* was modeled as a feedforward population that delivered synaptic input 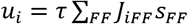 from a simulated population of Poisson processes of constant rate *r_FF_*. See Table 1 for network parameters.

**Table 1:** Parameters of the spiking network

|  |  |
| --- | --- |
| $N_E$, Number of excitatory cells | 1200 |
| $N_I$, Number of inhibitory cells | 300 |
| $N_{FF}$, number of feedforward cells | 2400 |
| $K_{EE}$, expected E->E connections per cell | 300 |
| $K_{IE}$, expected E->I connections per cell | 300 |
| $K_{EI}$, expected I->E connections per cell | 75 |
| $K_{II}$, expected I->I connections per cell | 75 |
| $K_{FF}$, number of feedforward connections per cell | 300 |
| $V_{th}$, threshold potential | 20 mV |
| $V_{rest}$, rest potential | 0 mV |
| $V_{reset}$, reset potential | 10 mV |
| $\tau_m$, neuronal time constant | 20 ms |
| $\tau_{ref}$, refractory period | 2 ms |
| $\tau_{delay}$, synaptic delay | [2-5ms] uniform distribution |
| $J_{FF}$, connection weight of FF connections | $(V_{th} - V_{rest})/\sqrt{K_{ff}}$ |
| $\tau_{STDP}$, plasticity time constant | 20ms |

iSTDP was implemented for inhibitory -> excitatory synapses as in (*27*), using a synaptic trace *x*, which increased to 1 at the time of spiking for inhibitory cells, increased to 1 following the synaptic delay for their postsynaptic excitatory partners, and decayed with time constant *τ_STDP_. J_ij_* for inhibitory->excitatory synapses *ij* followed the following plasticity rule:

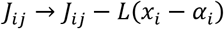

At every spike time of inhibitory cell *j*, following the synaptic delay and

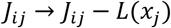

at the spike time of every excitatory cell *i*. As shown in (*27*), the effect of this rule is to adjust the inhibitory synaptic weights until each excitatory cell reaches its target rate, *α_i_* = 2*r_target_τ_STDP_*. Target rates were chosen from a lognormal distribution with *μ* = 0 and *σ* = 1.

## Suppplementary Figures

**Figure S1:**
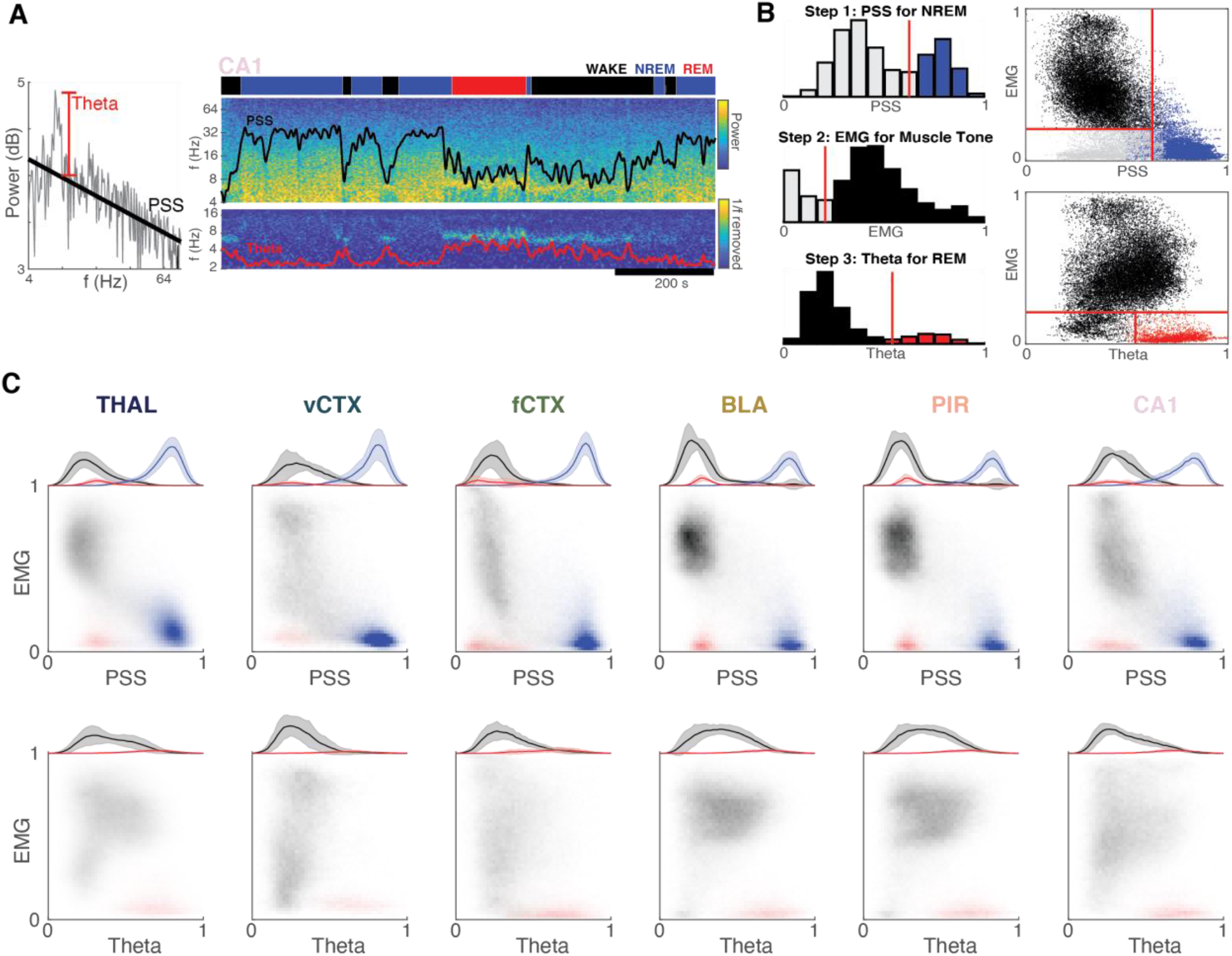
Brain state Scoring. **A**: State scoring metrics. Power spectrum slope (PSS) is the slope of a line fitted to the median-smoothed log-log power spectrum of the LFP, and estimates the 1/f exponent. Theta is taken to be the maximal power between 5-10Hz, after removal of PSS. Right: timeseries of PSS (black line) and theta power (red line) for a sample recording from the hippocampal CA1 pyramidal layer. Note low PSS values during theta states and high PSS values during NREM. **B**: State metric distributions for the example recording. Each point represents a time segment in the recording. States are clustered similar to (*14*), using a combination of thresholds in the distribution of PSS (Step 1), EMG (Step 2) and theta power (Step 3). Right top panel: PSS vs EMG for an example recording. Right bottom panel: Theta power vs EMG for the example recording. Red lines in each panel indicates the thresholds used to separate states. **C**: Mean state metric distribution (PSS, theta and EMG) over all recordings in each region. Brain state clustering is comparable across regions. The occupancy in each state depended on the animal’s behavior. In home cage recordings locomotion (wake; grey) and NREM (blue) dominate. Theta = normalized theta power

**Figure S2:**
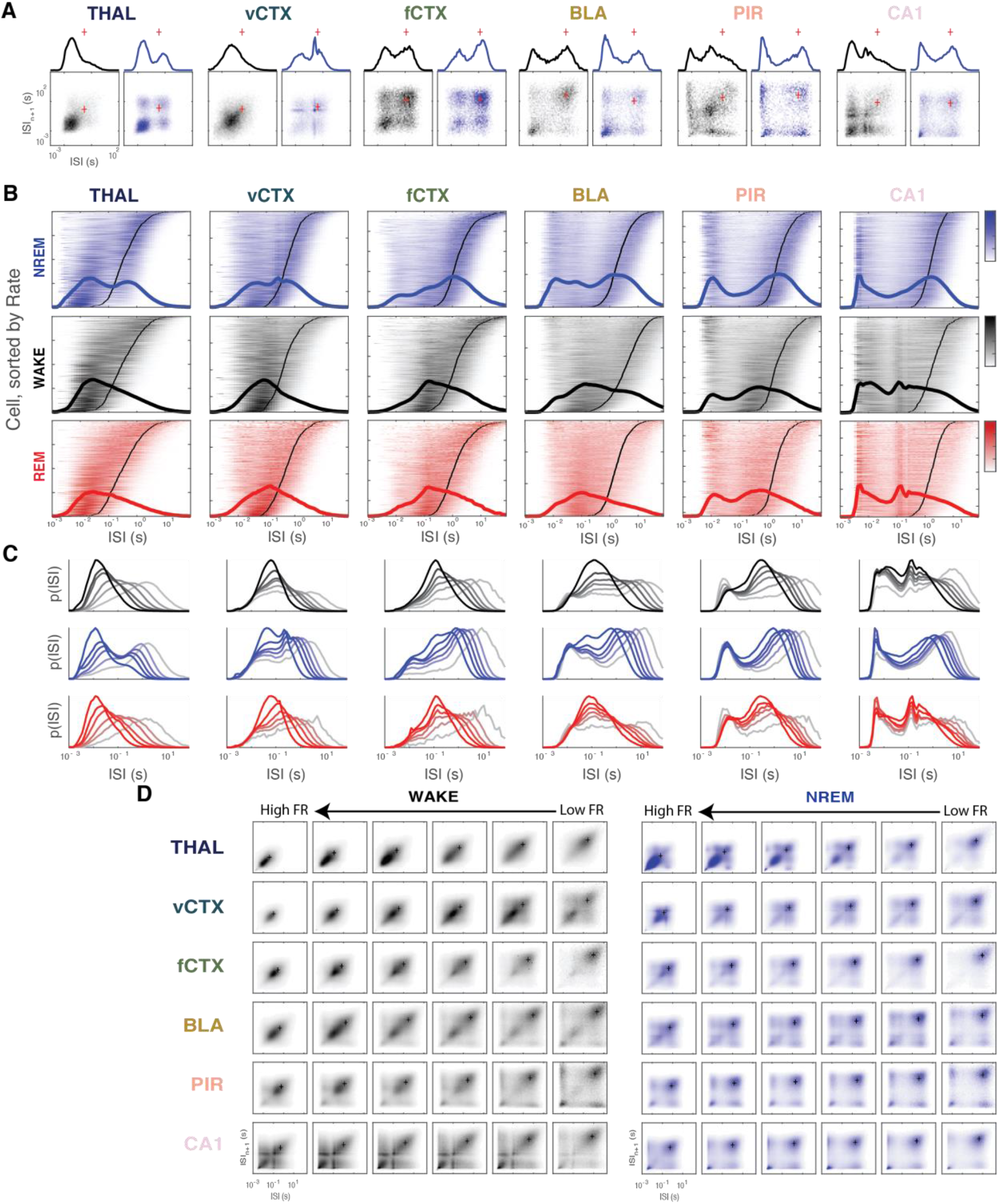
ISI distributions and return maps show region/state-specific spiking modes in forebrain excitatory neurons. **A**: ISI distributions and return maps for an example cell from each region, during WAKE (black) and NREM sleep (blue). Red cross indicates 1/mean firing rate. **B**: ISI distributions from all cells in the dataset, grouped by region and sorted by mean firing rate (black line: 1/mean firing rate). Thick lines indicate sum of all neurons. **C**: Mean ISI distributions shown separately for firing rate sextile groups in each region/state. **D**: Mean ISI return maps shown separately for firing rate sextile groups in each region/state. + marks 1/mean firing rate.

**Figure S3:**
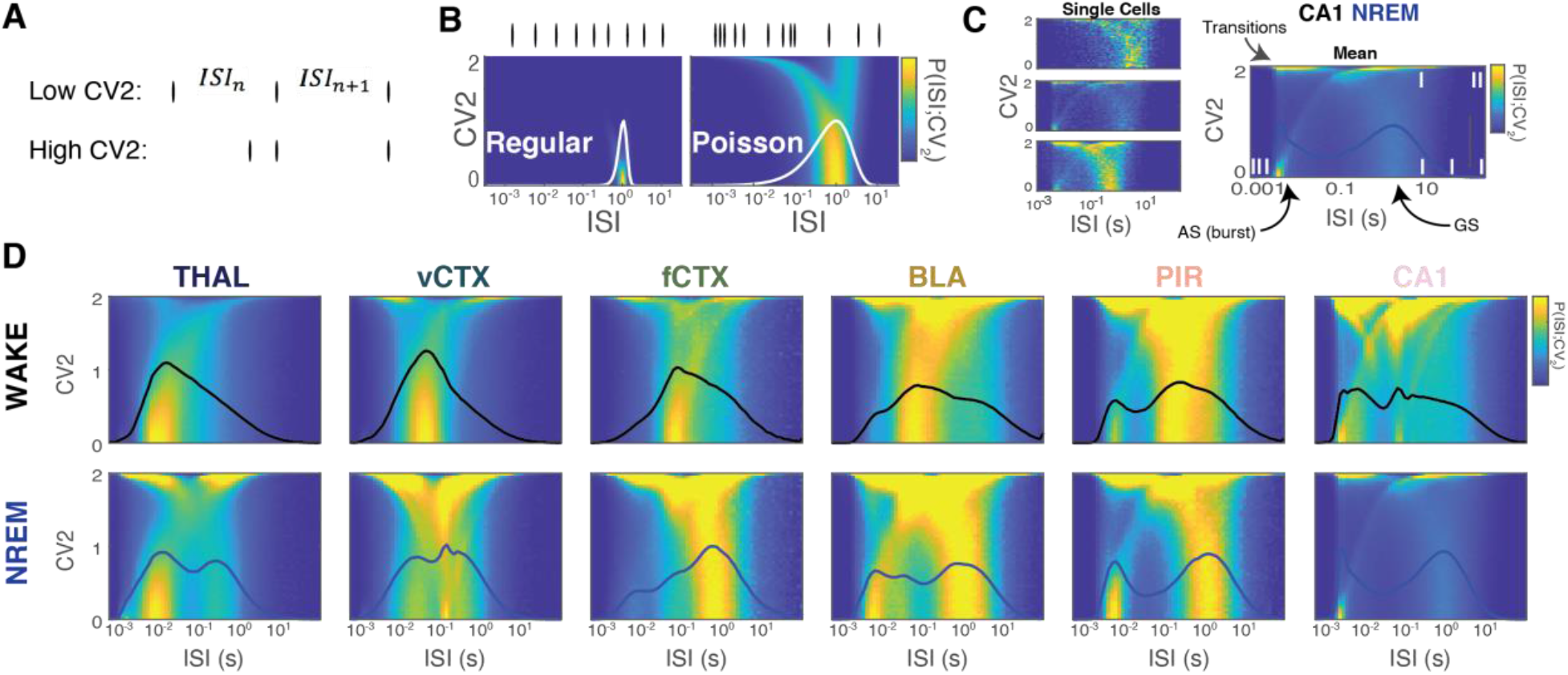
Timescale-specific spiking (ir)regularity. **A**: Coefficient of variation 2 (CV_2_), a local metric of spiking irregularity (*52*) **B**: Joint distribution of ISI and CV_2_ values (P(ISI;CV_2_)) for regular spiking and Poisson spiking model neuron. **C**: P(ISI;CV_2_) for three example CA1 cells during NREM sleep. (Right) mean P(ISI;CV_2_) over all CA1 cells during NREM sleep. **D**: Mean P(ISI;CV_2_) for each region during WAKE and NREM. Superimposed lines in all P(ISI;CV_2_) plots indicate mean ISI distribution.

**Figure S4:**
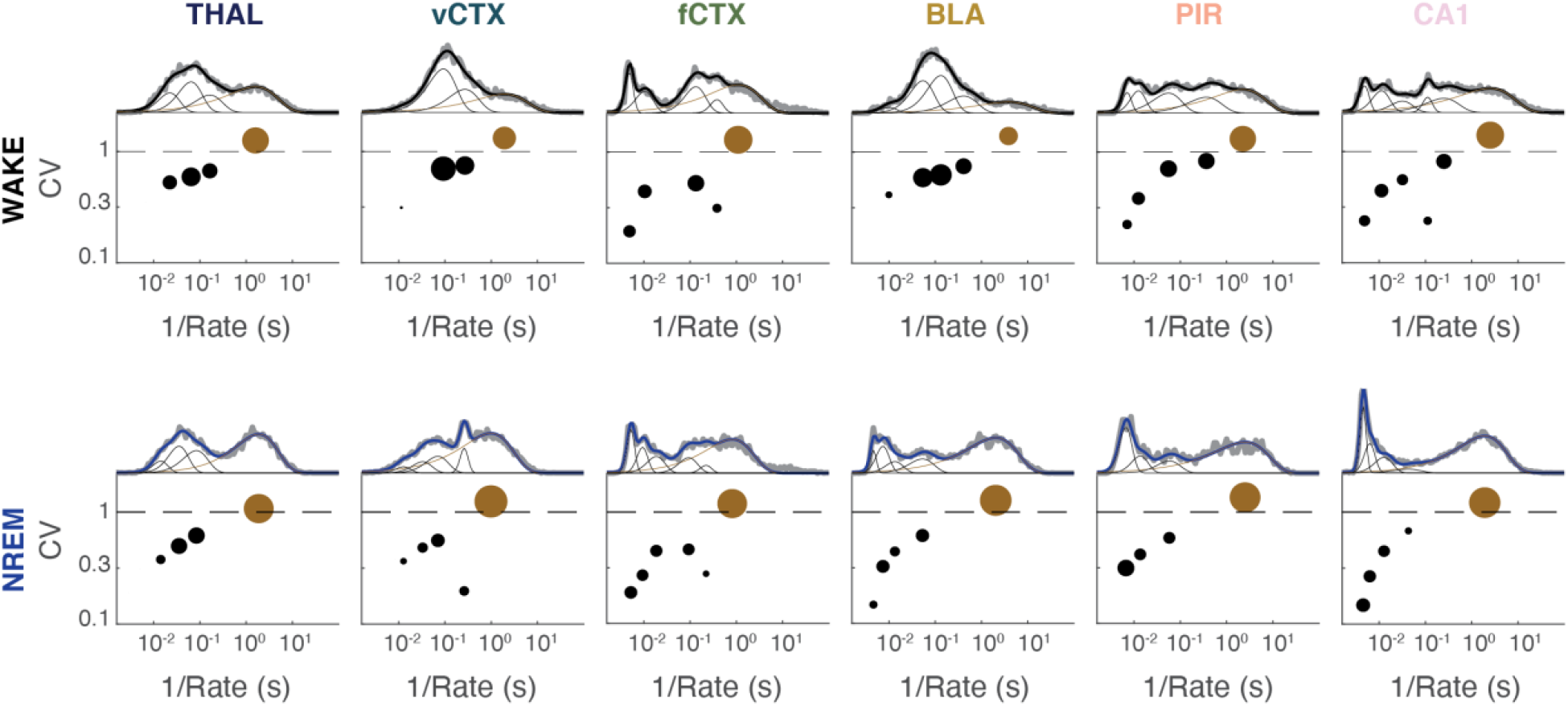
Mixture of Gammas model fits to example neurons. ISI distributions for an example cell from each region, during WAKE (top) and NREM sleep (bottom). Grey thick line is the observed ISI distribution, solid black line is the mixture of gammas (thin lines) fit, with each composite log-gamma distribution indicated as a point denoting its CV, Rate, and weight (black: AS modes, brown: GS mode). Weight is defined as the fraction of spikes in a given mode and its magnitude is proportional to the size of the circle.

**Figure S5:**
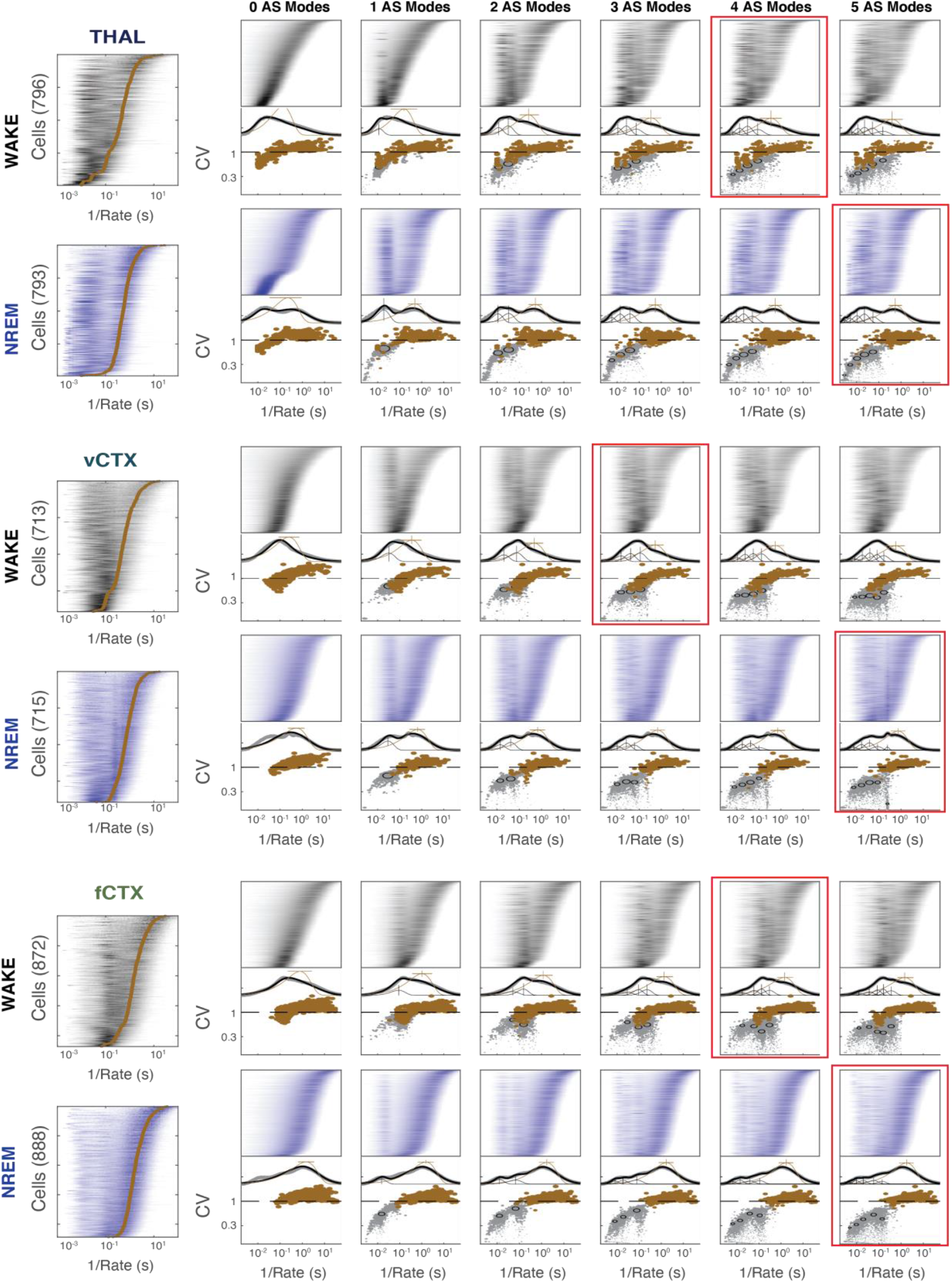
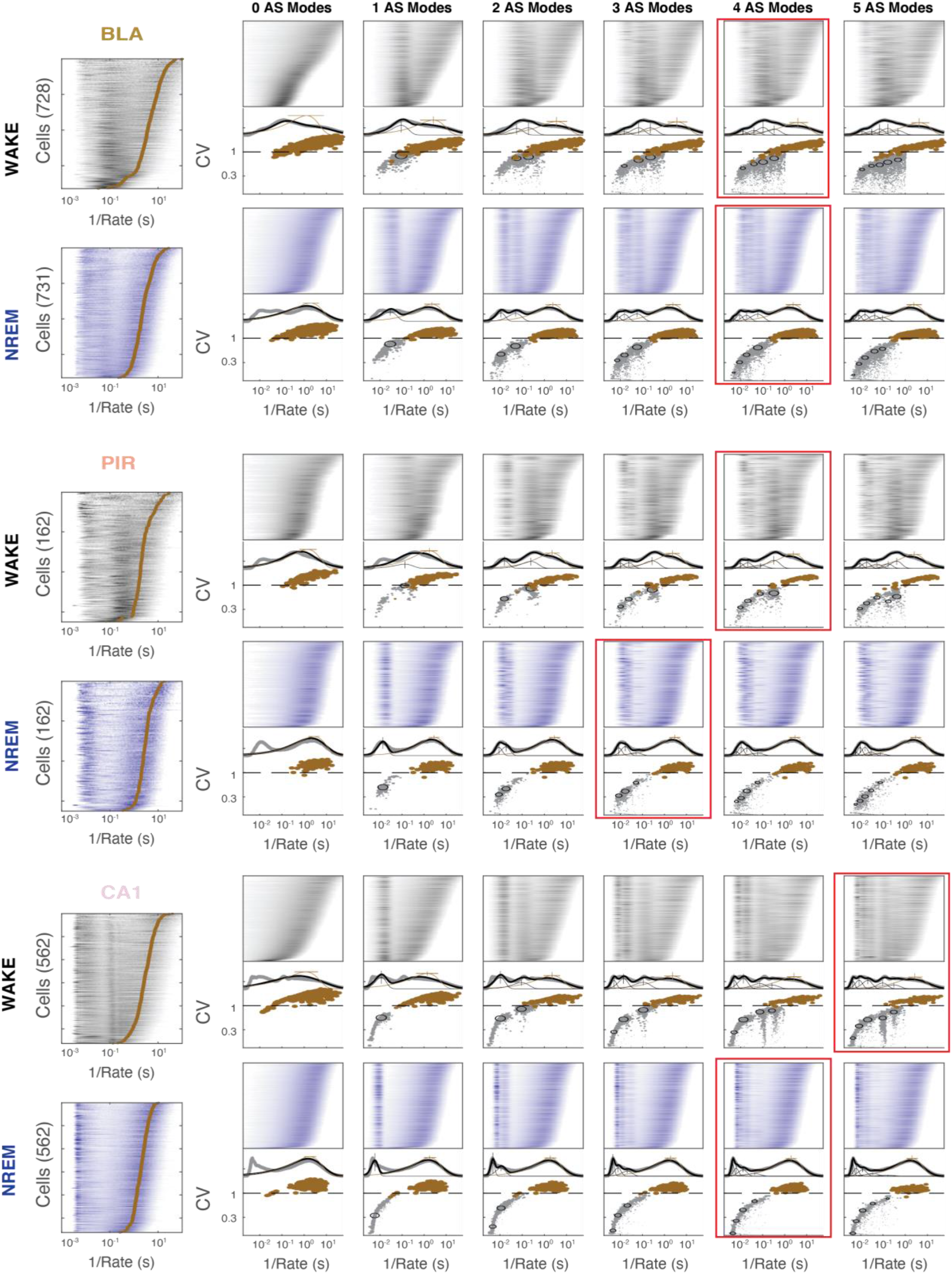
Mixture of Gammas model fits to all neurons. ISI distributions for all cells in WAKE and NREM state (left large panels). Right panels: ISI distributions of all cells and their fit to the mixture of gammas model (compare to Figure 2D) with 0 to 5 shared AS modes. Red square indicates the number of AS modes used for each region, determined via visual inspection of a number of heuristics: match between modeled and experimental ISI distributions, ability to capture known spiking patterns in each regions and ability to match mean ISI distribution. The lowest number of modes to fit these criteria was selected.

**Figure S6:**
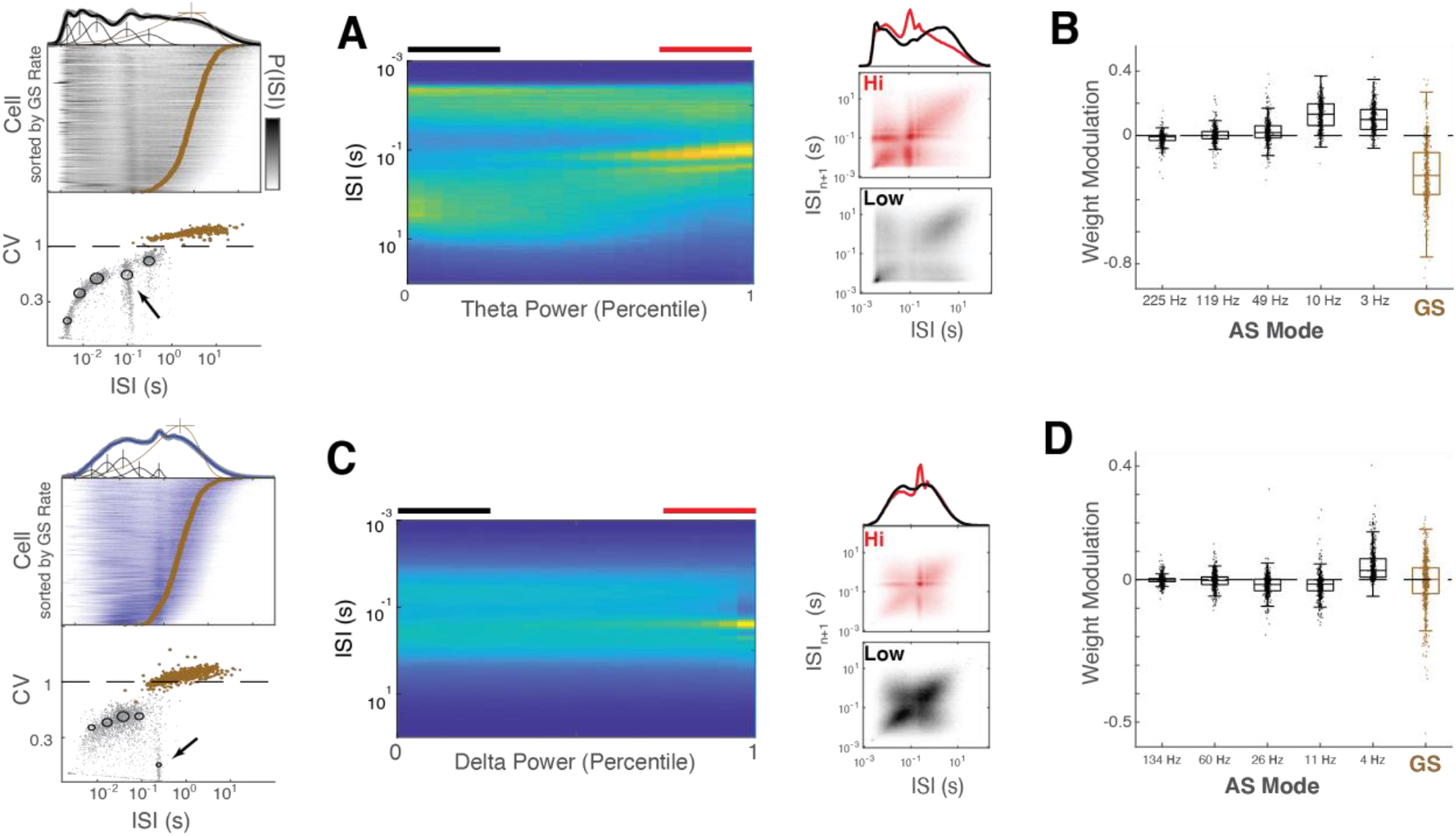
CA1 and vCTX AS spiking modes relate to theta and delta oscillations, respectively. **A:** ISI distribution conditioned on CA1 theta power during WAKE. Right: mean ISI distribution and return map during epochs of high and low theta power. Note similar spiking modes but different occupancies in high and low theta power epochs. **B:** Modulation of the weight of each spiking mode by the power of theta. Note largest modulation at 10 Hz, corresponding to theta band. **C:** ISI distribution conditioned on vCTX delta power during NREM. (Right) mean ISI distribution and return map during epochs of high and low delta power. **D:** Modulation of the weight of each spiking mode by the power of delta.

**Figure S7:**
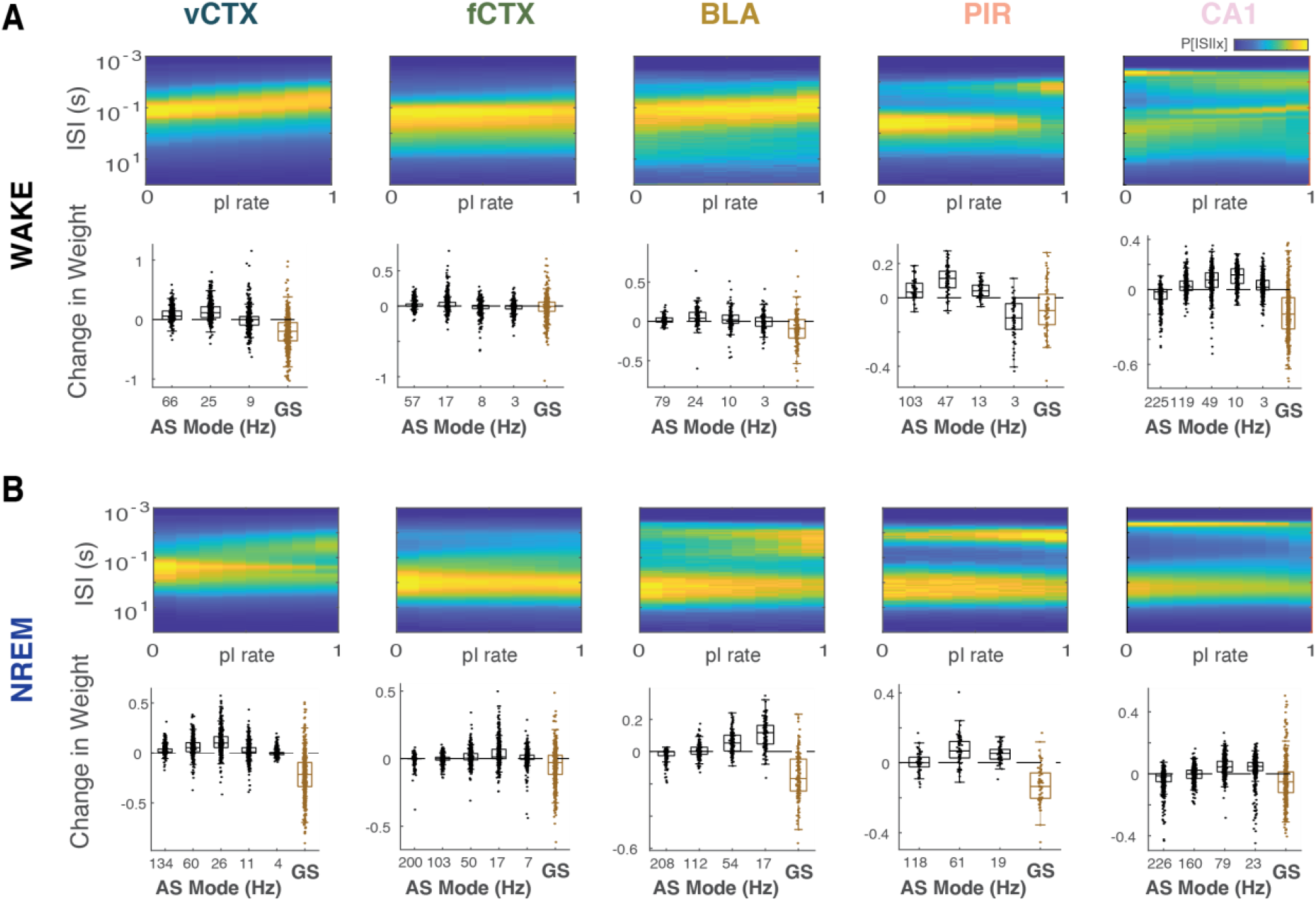
Interneuron activity and Spiking Modes. **A,B** Top panels: Mean ISI distribution conditioned by mean normalized rate of co-recorded interneurons during WAKE (A) and NREM (B) states. Bottom panels: Modulation of each ISI mode by interneuron rate, as quantified by the mixture of Gammas model.

**Figure S8:**
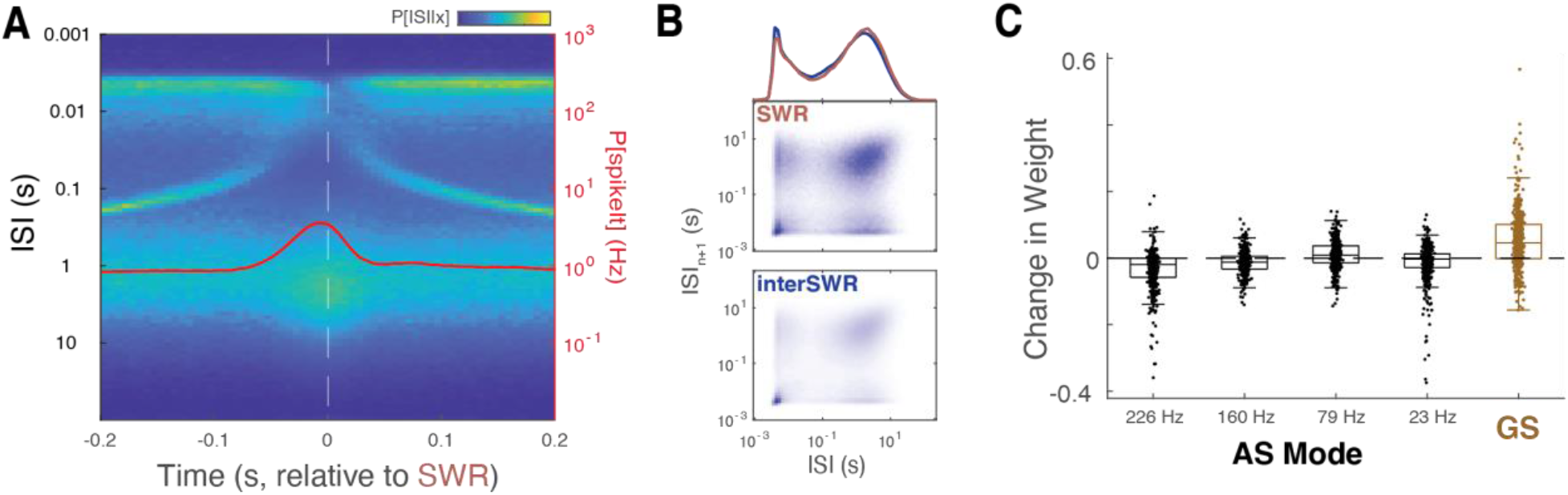
Spiking Modes During Hippocampal Sharp Wave Ripples (SPW-R). **A:** Mean peri-event ISI distribution, aligned to SPW-R peak (time 0). Note decreased conditional probability of spike bursts (< 6 ms ISI) during SPW-R. Red line, probability of spike occurrence of pyramidal cells. **B:** Mean ISI distribution and return map during SPW-Rs and inter-SPW-R periods. Note similar spiking modes but different occupancies during SPW-Rs and inter-SPW-R periods. **C:** Change in weight for each ISI mode between SPW-Rs and inter-SPW-R periods, as measured by the mixture of Gammas model for each CA1 cell.

**Figure S9:**
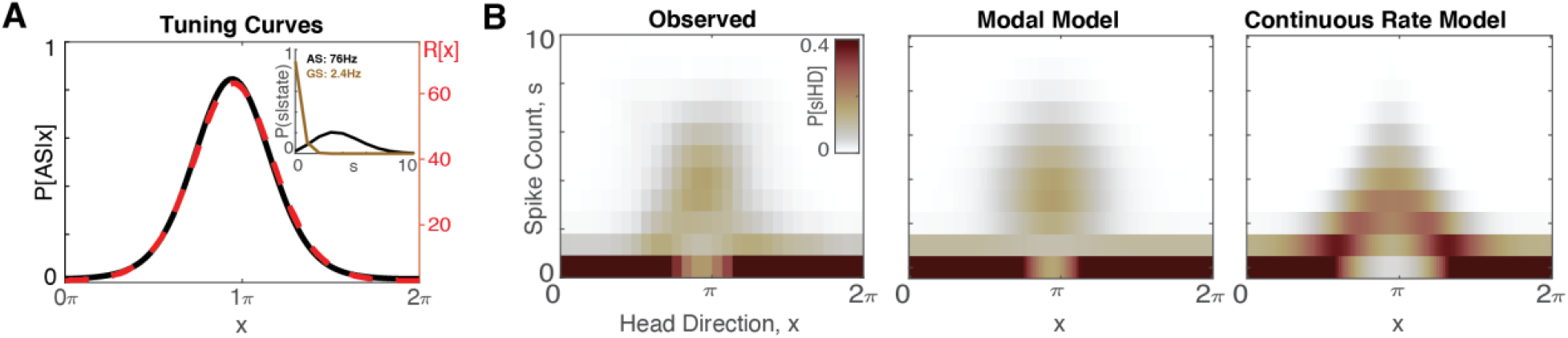
Modal Encoding Model for Head Direction Cells in Thalamus. **A: In** the continuous model (red dashed line), the tuning curve is the rate (right y axis) that varies continuously as a function of head direction (x). For the modal model (black line; left y axis), the tuning curve corresponds to the probability of the cell being in the activated state (AS). The inset shows the probability of observing a given number of spikes in a 50ms window when the cell is in GS (brown) and AS (black) mode. Thus, both models have a tuning curve that reflects the encoding of a stimulus variable in neural activity. In the continuous rate model, the stimulus is encoded by a continuous rate. In the modal model, the stimulus is encoded by occupancy in one of two discrete modes. Tuning curves fit is from the cell shown in B. **B:** (Left) Observed spike count given head direction for an example HD cell. (Middle and right) Spike count given head direction for the modal (middle) and continuous (right) model, with the best-fitting parameters for the example cell.

**Figure S10:**
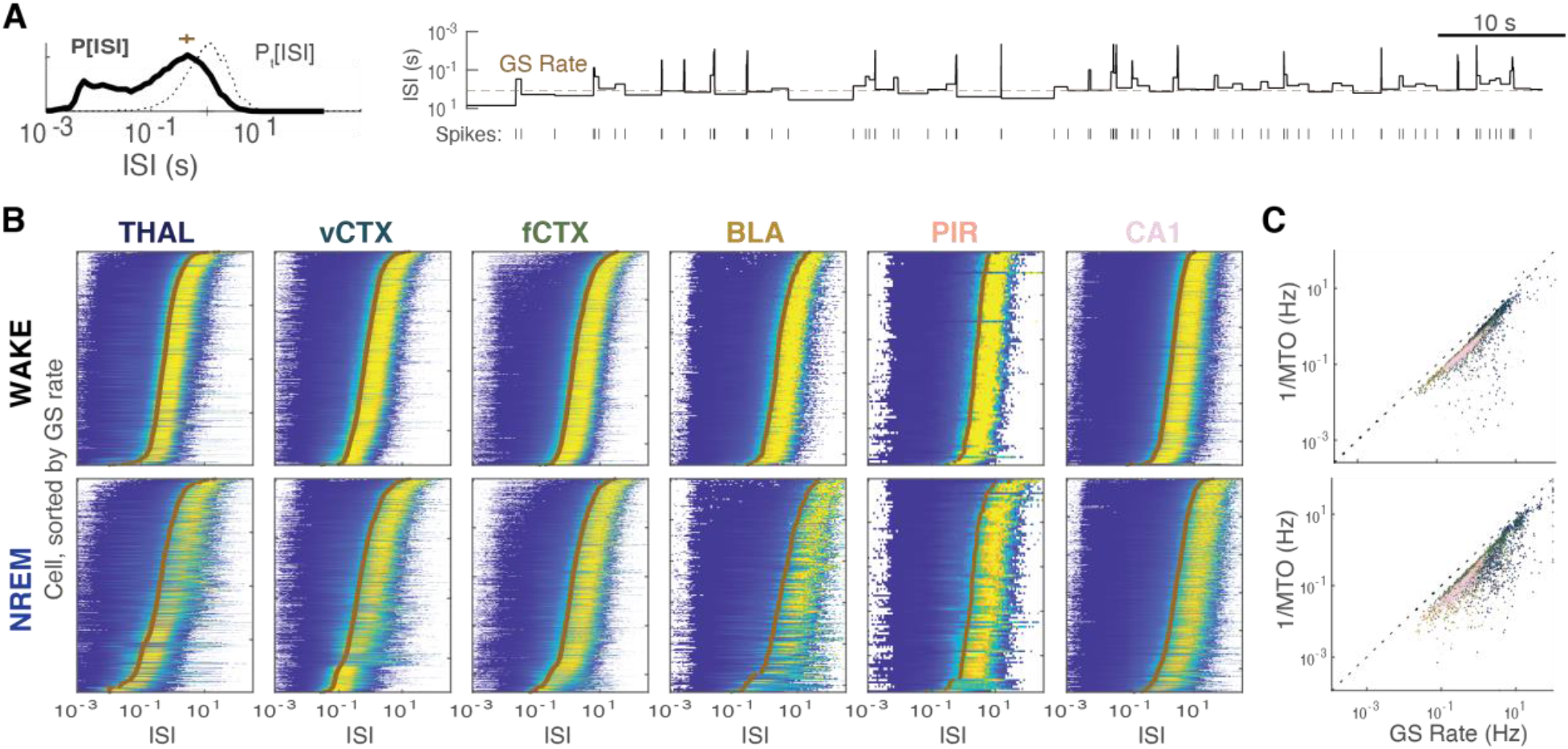
Neurons spend the majority of time in GS mode. **A**: ISI distribution (black line) and time occupancy distribution (dotted line), defined as the percentage of time within a given ISI, for an example CA1 cell during NREM sleep. Right panel: Time series of ISIs during an example 100s of recording time. Brown dashed line: GS rate. **B**: Time occupancy distribution, for all cells in each region/state. Cells were sorted by GS rate (brown) **C**: GS rate vs medium time occupancy (MTO) for all cells during WAKE and NREM states. Dot colors correspond to the respective brain regions (as in B).

**Figure S11:**
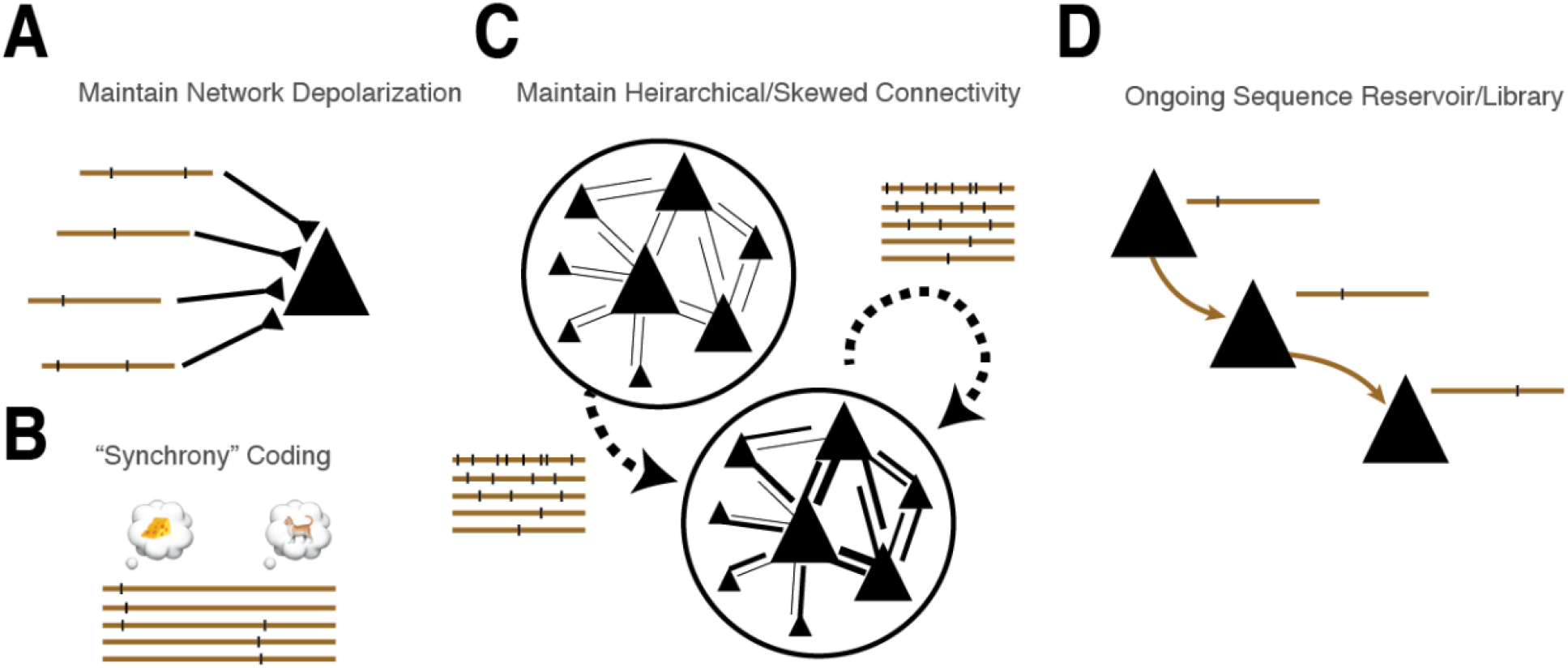
Hypothesized Functions of GS. **A**: GS activity maintains network excitability, ongoing/spontaneous activity, and sensitivity to perturbation. **B**: GS spikes represent sensory/motor/cognitive variables through synchrony and cell identity. **C**: Heterogeneous balance maintains a hierarchical synaptic network. **D**: GS spikes act as a “reservoir” or library of sequences that can be selected to correspond to input patterns. Mechanisms of irregular activity in cortical neurons in vivo have been the subject of extensive study. Irregular spiking is thought to come from a regime in which the balance of excitatory and inhibitory synaptic inputs keeps the membrane potential below the threshold for spike generation, and spikes are fluctuation-driven due to the balance of excitatory and inhibitory inputs (*2*, *4*, *53*), which can emerge under a range of conditions in recurrent (*7*) and feedforward (*54*) networks. GS mode reflects isolated spikes of single neurons that may result from the combination of ongoing network-induced fluctuations in membrane potential and intrinsic properties of the neuron. We found that a self-balancing network is able to reproduce GS-like activity, favoring the role of a network effect. The conservation of a neuron’s GS rate across brain states (i.e., Fig. 4F) and the independence of GS spikes across neurons suggest that GS rate is a characteristic property of each neuron (“fingerprint”;(*55*)). However, an open question is the source of GS rate heterogeneity – why are some neurons low rate in the GS (e.g., < 0.01Hz in some neurons), while others have faster GS spikes with a distribution of at least two orders of magnitude? In our model, we used an inhibitory spike time dependent plasticity (iSTDP) rule by which each neuron homeostatically adjusts the strength of its incoming inhibitory synapses to reach a unique specified target rate (*27*), resulting in a GS rate for each cell set by the strength of incoming inhibitory synapses relative to excitation. However, a number of other homeostatic processes might be able to achieve the same outcome (*55*–*57*), including intrinsic excitability, Na^+^ channel recovery (*58*, *59*), excitatory synaptic, adaptation (*60*), resting/equilibrium voltage, or other Ca^2+^ - mediated processes. Alternatively, heterogeneity in GS rate might be an emergent property of a network (*61*) or its development, and may not need to be regulated by a cell-autonomous homeostat. Given the central importance of maintaining a sustained network dynamic, it is likely that several mechanisms contribute to the generation of GS activity. Further work is needed to identify specific signatures of these putative sources of heterogeneity, and to compare their implications for network function.

